# Re-sensitising XDR biofilms using a combination of bacteriophage cocktails and colistin: a natural approach

**DOI:** 10.1101/2022.02.04.479063

**Authors:** Naveen Kumar Devanga Ragupathi, Dhiviya Prabaa Muthuirulandi Sethuvel, Mohanraj Gopikrishnan, Dhivya Murugan, Ramya Juliet, Monalisa Majhi, Malathi Murugesan, George Priya Doss C, Leshan Wannigama, Peter N. Monk, Esther Karunakaran, Balaji Veeraraghavan

## Abstract

Persistent antibiotic use results in the rise of antimicrobial resistance with limited or no choice for multidrug resistant (MDR) and extensively drug resistant (XDR) bacteria. This necessitates a need for alternative therapy to effectively combat clinical pathogens that are resistant to last resort antibiotics. The study investigates hospital sewage as a potential source of bacteriophages to control MDR/XDR bacterial pathogens. 81 samples were screened for phages against selected clinical pathogens. 10 phages were isolated against *A. baumannii*, 5 phages against *K. pneumoniae* and 16 phages obtained against *P. aeruginosa*. The novel phages were observed to be strain-specific with a complete growth inhibition of up to 6 hrs. Phage plus colistin combinations further reduced the MBEC of colistin up to 16 folds. Notably, cocktail of phages exhibited supreme efficacy with complete killing at 0.5-1 µg/ml colistin concentrations. Thus, phages specific to clinical strains has a higher edge in treating nosocomial pathogens with their proven anti-biofilm efficacy. In addition, analysis of phage genomes revealed close phylogenetic relations with phages reported from Europe, China and other neighbouring countries. This study serves as a reference and can be extended to other antibiotics and phage types to assess optimum synergistic combinations to combat various drug resistant pathogens in the ongoing AMR crisis.

## Introduction

Persistent use of antibiotics has led to the emergence of multidrug resistant (MDR) and extensively drug resistant (XDR) bacteria, making the most effective drugs largely ineffective. Consequently, carbapenem and colistin resistant Gram-negative bacteria have emerged as an important therapeutic challenge. Development of novel antibiotics to treat drug resistant nosocomial infections, especially those caused by ESKAPE pathogens (*Acinetobacter baumannii, Klebsiella pneumoniae*, and *Pseudomonas aeruginosa*), is an urgent need (Mulani et al., 2019). The formation of biofilm by these pathogens provides further challenges to treatment due to the enhanced survival of the organisms on hospital equipment, increased antibiotic resistance, virulence, and re-occurrence (Vukotic et al., 2020).

Biofilms are a collective of single or multiple types of microorganisms that can grow on many abiotic and biotic surfaces. These adherent cells are embedded within a self-produced matrix of extracellular polymeric substances such as polysaccharides and e-DNA. The Centers for Disease Control estimates that over 65% of nosocomial infections are caused by biofilms. Available antibiotics and common disinfectants have shown limited ability to remove biofilms, due largely to reduced penetrance and altered microbial metabolism (Sharma et al., 2019). Recently, bacteriophage (phage)-based treatments involving a single phage or phage cocktails, phage-derived enzymes, and phage in combination with antibiotics have become interesting alternatives for biofilm removal.

Among the ESKAPE pathogens, *A. baumannii, K. pneumoniae*, and *P. aeruginosa* are the leading cause of nosocomial infections amongst critically ill and immunocompromised individuals; these pathogens harbour a variety of drug resistance mechanisms (Santajit and Indrawattana, 2016). *K. pneumoniae* has become a relevant healthcare-associated pathogen, causing approximately 14–20% of the infections related to respiratory tract, lower biliary duct, surgical wounds, and urinary tract infections. Molecular strategies to develop carbapenems resistance are carbapenemases production (*bla*_KPCs_, *bla*_OXA_ and metallo-β-lactamases); non-expression or mutation of porin genes (OmpK35 and OmpK36); upregulation of efflux systems.

*A. baumannii* is a ubiquitous pathogen capable of causing both community and health care associated infections, including septicemia, bacteremia, ventilator-associated pneumonia, wound sepsis and endocarditis. According to CDC, 80% of *A. baumannii* is found primarily in hospitals and poses a risk to people who have suppressed immunity. A diverse array of transposons, plasmids, and insertion sequence elements along with antimicrobial resistance genes were reported to confer resistance to multiple antimicrobials (Rozwandowicz et al., 2018; Razavi et al., 2020). Currently no antibiotic treatment regimens work for XDR *Acinetobacter* infections. Similarly, *P. aeruginosa* mainly infects immunocompromised patients with cystic fibrosis, severe burns, cancer, and AIDS. Although the import of resistance mechanisms on mobile genetic elements is always a concern, the most difficult challenge with *P. aeruginosa* is the chromosomally encoded AmpC cephalosporinase, the outer membrane porin OprD, and the multidrug efflux pumps.

Treatment options for these pathogens are limited due to the emergence of XDR/PDR strains worldwide. Thus, there is an ongoing discussion as to whether phages can be an alternative to combat drug resistant bacteria. Assuming that there is no negative interference, the benefit of a simultaneous use of phage with antibiotic can be two-fold: In fact, cross-resistance between an antibiotic and phage seems less common than between two different types of antibiotics or phages (Jansen et al., 2018). In addition, unlike broad-spectrum antibiotics, phages are highly specific, and they do not kill the commensal microbiota, which are vital for patients with malnutrition and immunodeficiency (Nagel et al., 2016).

Phages are widely found in diverse environments, sewage effluent, soil, water, fecal materials, as well as the gastrointestinal tract of humans and animals; and they can be rapidly isolated (Pereira et al., 2017). In this study, we isolated phages from hospital sewage against drug resistant biofilm producing *K. pneumoniae, A. baumannii* and *P. aeruginosa* as target hosts. We investigated phage host range and stability along with efficacy against both planktonic and biofilm-embedded bacterial cells, by determining minimum inhibitory concentration (MIC), and minimum biofilm eradicating concentration (MBEC) of phage and phage-antibiotic combination. In addition, genetic relatedness of the isolated phages was studied using phylogenetic analysis.

## Methods

### Isolates

Clinical multi-drug resistant (MDR) isolates of *A. baumannii* (*n* = 15), *P. aeruginosa* (*n* = 8) and *K. pneumoniae* (*n* = 5) were collected from bloodstream infections and respiratory infections from patients admitted in a tertiary care hospital between 2018 to 2020. Isolates were identified and characterised using standard biochemical tests.

### Minimum Inhibitory Concentration (MIC)

MIC values of colistin were estimated by broth microdilution assay as per CLSI M100-S30, 2020 and interpreted accordingly (CLSI, 2020). Quality control strains *mcr-1* positive *E. coli* with 4 – 8 µg/ml expected MIC range, *E. coli* ATCC 25922 with 0.25 – 2 µg/ml and *P. aeruginosa* ATCC 27853 with 0.5 – 4 µg/ml were employed in all the test batches.

### Sewage sample collection

Sewage water was collected as single-point collections from 5 hospital sewage sites in and around Vellore (CMC 12.925743°N, 79.133804°E; Puliyanthangal 12.964527°N, 79.292959°E; Sripuram 12.870004°N, 79.090083°E; Vellore 12.915576°N, 79.133345°E; Ariyur 12.873067°N, 79.102310°E). Samples were collected in 500 ml plastic sterile bottles. Primary filtration was carried at the site of collection using a gauze cloth, to remove particulate matter. In the laboratory, 250 ml of sewage (both post & pre-treated) was re-filtered using Whatman Grade 1 filter paper and stored with the remaining 250 ml at 4 °C. Peter Monk overnight cultures of the target bacteria (host), adjusted to 0.5 McFarland, were added l to 150 ml of Tryptic Soy Broth (TSB) and incubated at 37°C for 1 hr in a conical flask. After incubation, 200 ml of filtered sewage was added to culture flasks and incubated overnight at 37°C, 120 rpm in a shaker incubator. Following incubation, the contents of the conical flasks were transferred to 50 ml tubes and centrifuged at 11,200 x g for 10 min. Supernatant was collected and filtered using Whatman filter paper. 2 ml of filtered supernatant was taken and centrifuged again at 11,200 x g for 10 min to collect clear supernatant. This was further subjected to filtration through a 0.22 µm syringe filter and utilized for further applications in the study.

#### Spot assay for phage screening

Method followed was as described by Assafiri et al. (2021) with slight modifications. Briefly, 100 µl of bacterial culture at log phase was spread on LB agar and allowed to dry for 10 min at room temperature (RT). 10 µl of phage suspensions obtained from processed sewage samples were spotted on plates with respective host target bacteria. Plates were allowed to dry in RT for 10 min, followed by incubation at 37 °C for 24 hr. Phage suspensions forming clear zones were recorded as harbouring bacteriophages specific to the target host.

#### Double agar overlay assay

Double agar overlay assay was performed for the screened phage suspensions as described earlier (Assafiri et al., 2021). 500 µl of the phage suspension and 200 µl of the 0.5 McFarland Standard adjusted target bacterial cultures were added to 4 ml of soft agar (0.4%). Host-phage interaction was facilitated by incubating at RT for 10 min. Prepared solution was poured forming a soft agar layer over the base agar layer (1%). Plates were left to settle and dry for 20 min at RT. Plates were further incubated at 37 °C overnight and formation of Bull’s eye plaques were monitored.

### Phage antimicrobial susceptibility testing (MIC)

MIC testing for phages was performed by broth microdilution method. *E. coli* ATCC 25922 strain was used as a quality control strain. A time-dependent approach was employed with different incubation times of 3, 6, 9, 24, 27, 30 and 48 hr to analyse the growth inhibition capacity of the phages and phage + colistin (sub-MIC 0.5 and 0.25x) against clinical strains. Results were interpreted based on the visual clearance of wells viewed against a black background.

### Phage stability analysis

A thermal stability assay was employed to evaluate the heat resistance of isolated phages. Phage suspensions (SM buffer: mixture of Sodium chloride, Magnesium sulphate and gelatin) were incubated at different temperatures (4 °C, 25 °C, 37 °C, 50 °C, 60 °C, and 70 °C) for 1 hr and the stability was measured by a soft agar overlay method. To assess pH stability, the phage suspensions were preincubated at pH 3-13 at 37 °C for 1 hr. After the incubation period, the phage titer was determined by double agar layer method. To evaluate the phage stability under UV light, the phage suspension was kept at a different time interval of 5 secs to 2 min, adapted and modified from (Mallick et al., 2021). Subsequently, the survival percentage of bacteriophage was measured by the soft agar overlay method.

### Biofilm screening of the hosts

A previously described method was used with minor modifications (Di Domenico et al., 2016; Devanga Ragupathi et al., 2020). Briefly, 150 µl of 0.05 OD_625_ culture was inoculated in a “96 well” microtiter plate and incubated at 37 °C for 24 hr. Biofilms were washed with water and stained with 200 μl of aqueous 0.1% (w/v) crystal violet. Wells were washed after 10 min and destained using 200 μl of 33% (v/v) glacial acetic acid after 5 min incubation at RT and OD_570_ was measured. Assays were performed in triplicates and a blank well containing only medium was used as negative control.

The OD_570_ values were used as a semi-quantitative biofilm production. Accordingly, the cut-off OD (ODc) was calculated as the mean OD of the negative control with three standard deviations. Biofilm production was classified as: OD < ODc = poor biofilm producer; ODc < OD ≤ 2 × ODc = weak biofilm producer; 2 × ODc < OD < 4 × ODc = moderate biofilm producer; and OD ≥ 4 × ODc = strong biofilm producer.

### Minimum biofilm eradication concentration (MBEC) for phages

#### Colistin monotherapy (CMT) and combinations

*The* MBEC assay was performed as previously described (Chen P. et al., 2014) with slight modifications (Devanga Ragupathi et al., 2020). Briefly, all isolates were grown at 37 °C overnight in cation-adjusted MHB with 1% glucose. Cultures were then adjusted to 0.05 OD_600_ with MHB and 150 μL of this adjusted inoculum was added to all the wells (except one well used for sterility control) in the 96-well MBEC Assay^®^ plate (Innovotech, AB, Canada). The MBEC inoculator plate was then inserted and incubated at 37 °C for 24 hr to allow biofilm formation. After the biofilm formation in the inoculator plate, four pegs were collected aseptically and tested for the density of the formed biofilm.

The removed pegs were put in a fresh 96-well plate (A1–A4) with 200 μl TSB medium with 1% Tween20 (rich medium) and sonicated in high power for 10 min. After sonication, 20 μl of the inoculum from the suspension (planktonic cells) in the MBEC bottom plate (any four random wells) were inoculated in four wells (A5–A8) with 180 μl of rich medium. The suspensions were then serially diluted up to 10^−8^ dilutions, and 10 μl was plated on LB agar followed by incubation at 37 °C overnight for measuring cell count (CFU/ml).

The rest of the inoculator plate with biofilm was inoculated into a 96-well plate containing various concentrations of the antibiotic in MHB (200 μl) in duplicates. The MBEC plate setup with antibiotics was incubated at 37 °C for 24 hr. The inoculator plate was removed and washed with sterile distilled water for 1 min and introduced in a 96-well bottom plate with 200 μl of rich medium. The plate was sonicated for 10 min at high power. Released biofilm cells (sonicated) and planktonic cells from MBEC bottom plate were serially diluted up to 10^−4^ dilutions and the cell counts were determined as above to enumerate MBEC values.

### Genome sequencing

#### Phage DNA extraction

Phage DNA was extracted from 1 ml of fresh lysate using a modified phenol:chloroform method (Clokie et al., 2003). Briefly, cell debris was pelleted by centrifugation (13,000× *g* for 10 min) at 4 °C. The supernatant was collected, transferred to a fresh tube, and the process repeated. The final supernatant was mixed with an equal volume of phenol (pH 10), vortexed for 30 s and centrifuged at 13,000× *g* for 10 min at 4 °C. The aqueous layer was mixed with an equal volume of phenol:chloroform (1:1), vortexed for 30 sec, and re-centrifuged at 13,000× *g* for 10 min at 4 °C. Finally, the aqueous layer was extracted, mixed with an equal volume of phenol:chloroform:isoamylalcohol (25:24:1), and the process was repeated again. The aqueous layer was extracted again, mixed with 1/10th volume 7.5 M ammonium acetate, two volumes of ice cold 100% ethanol and the DNA pellet was collected by centrifugation at 13,000× *g* for 30 min at 4 °C. The pellet was washed twice in 70% ethanol, dried, and resuspended in nuclease-free water prior to quantification with Qubit (Life Technologies). DNA was diluted to 0.2 ng/μl and libraries were prepared using the NexteraXT (Illumina) protocol following the manufacturer’s instructions.

#### Bacterial host DNA extraction

Genomic DNA of host isolates were extracted with QIAamp DNA mini kit (Qiagen, Hilden, Germany). Host isolates were subjected to sequencing to analyse the prophage regions and genetic factors responsible for antimicrobial resistance and virulence/toxins.

#### Bacterial whole genome and phage sequencing

Both phage and host genomes were sequenced using Illumina iSeq100 according to the manufacturer’s instructions. The data was assembled *de novo* using Unicycler v0.4.8. The sequence was annotated in prokka v1.14.6 (Seemann et al., 2014) and NCBI Prokaryotic Genomes Automatic Annotation Pipeline (PGAAP) (https://www.ncbi.nlm.nih.gov/genome/annotation_prok/).

### Bacterial host genome analysis

Assembled sequences were analysed using ResFinder and PlasmidFinder from the Center for Genomic Epidemiology (CGE) server (http://www.genomicepidemiology.org/services/). MLST of the isolates was identified using MLST 2.0 (https://cge.cbs.dtu.dk/services/MLST/). Prophage regions were identified using PHASTER (Arndt et al., 2016).

### Phage genome analysis

Assembled fasta sequences were analysed using PHASTER to identify the phage regions. Phage sequences were then extracted and analysed further. Easyfig v2.2.2 was used to compare the phage sequences to identify common regions (Sullivan et al., 2011). Phages from the study were compared with global sequences to develop a SNP phylogeny for closely related phages. The phylogenetic tree was visualised in iTOL v6.

### Scanning electron microscopy (SEM) of phages

Biofilms were grown on coverslips using culture conditions as described above. SEM analysis was performed as mentioned earlier (Dassanayake et al., 2020). Collagen coating (2 mg/ml) was used to support adhesion of bacterial cells on glass surfaces. Biofilms on glass disks were fixed in 2.5% phosphate-buffered glutaraldehyde for 1 hr at 4 °C and dehydrated in a graded series of cold ethanol-water mixtures (10, 20, 30, 50, 70, 80, 90, 95, and 100% ethanol) for 10 min each. Samples were dehydrated by critical point drying and coated with palladium-gold using a Hummer 6.2 sputter coater. After processing, samples were analysed in an Evo 18 research scanning electron microscope (ZEISS Research Microscopy Solutions, Jena, Germany) in high vacuum mode at 2 kV.

### Statistical Analysis

Clinical characteristics of infections were recorded in Microsoft Excel 2016 (Roselle, IL, United States) and analyzed for significance in SPSS 16.0 using *t*-test. Logarithmic growth values of antimicrobial treated biofilms were calculated as mean ± SD of triplicate values. GraphPad Prism v8.2.0 was used to generate the MBEC graphs.

## Results

### Characterisation of clinical isolates (host)

A total of 25 isolates were selected as hosts for screening of phages. This includes *A. baumannii* (*n* = 12), *P. aeruginosa* (*n* = 8) and *K. pneumoniae* (*n* = 5). Clinical outcome, along with comorbidities and antibiotic intake of these isolates, are listed in Table 1.

**Table 1:**
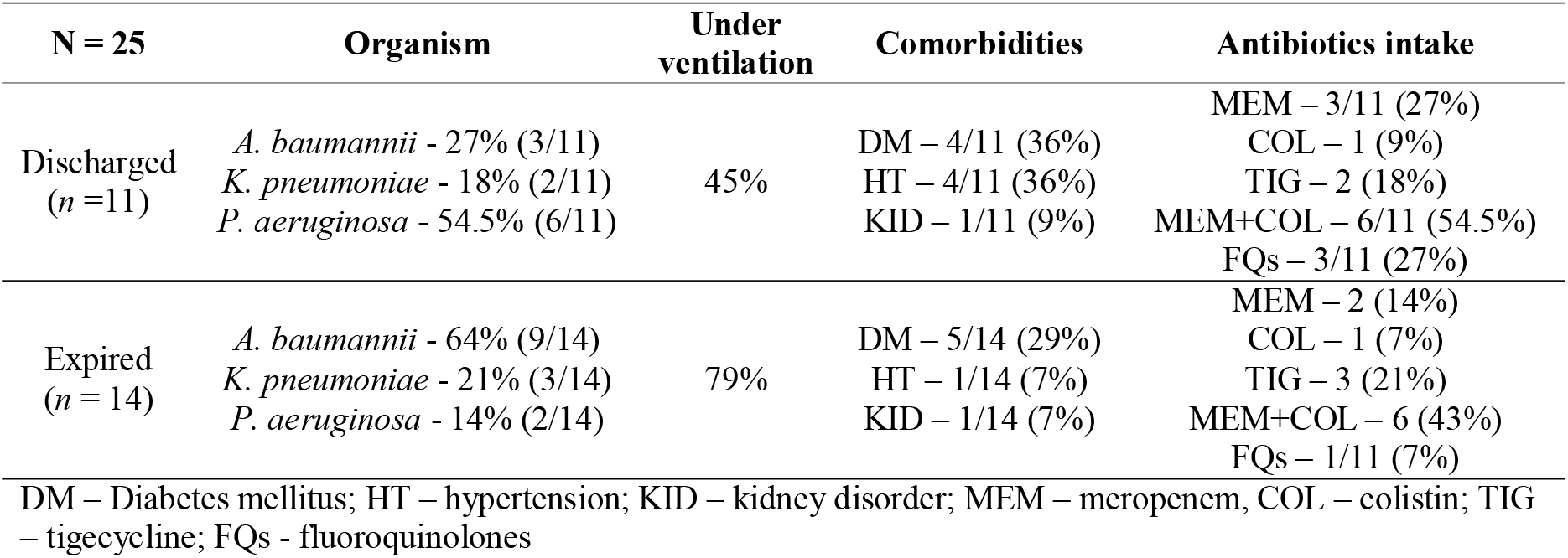
Clinical details of the Gram-negative isolates selected for screening of bacteriophages

Ventilation is one of the risk factors for mortality which was higher in patients with *A. baumannii* and *K. pneumoniae* when compared with *P. aeruginosa*. There was no correlation with comorbidities and mortality, which shows that the other mentioned factors were the drivers of mortality. All other clinical details related to the selected host strains are given in Table S1.

### Biofilm screening of host

The bacterial host isolates were analysed for their ability to form biofilms *in vitro*. All isolates were strongly biofilm forming with >4 fold higher than the control. Accordingly, the capacity of hosts to form biofilm ranged from 4.8 to 13.3 folds for *A. baumannii*, 5 to 6.5 for *K. pneumoniae* and 9.8 to 11.7 for *P. aeruginosa*.

### Isolation of bacteriophages and their spectrum of activity against clinical strains

A total of 81 sewage samples were screened against clinical strains of *A. baumannii* (*n* = 12), *K. pneumoniae* (*n* = 5), and *P. aeruginosa* (*n* = 8). Of these 81 samples, 10 phages were isolated against *A. baumannii* (*n* = 6); among these, 5 phages were specific to one or two strains, while the remaining 5 phages were active against >4 strains. For *K. pneumoniae* (n = 3), 5 active phages were isolated, with 1 having broad spectrum activity on 3 strains, while remaining 4 phages exhibited activity against two or three strains. For *P. aeruginosa* (*n* = 4), 16 phages were isolated, of which, 6 phages were active against 3 strains, and the remaining 10 strains were active against at least 2 strains. All the 31 isolated phages were labelled with the prefix BNDM consecutively. Double agar overlay assays for all 31 phages were performed against the specific bacterial hosts, which revealed the phenotypic characteristics of the phages as given in Table 2.

**Table 2:**
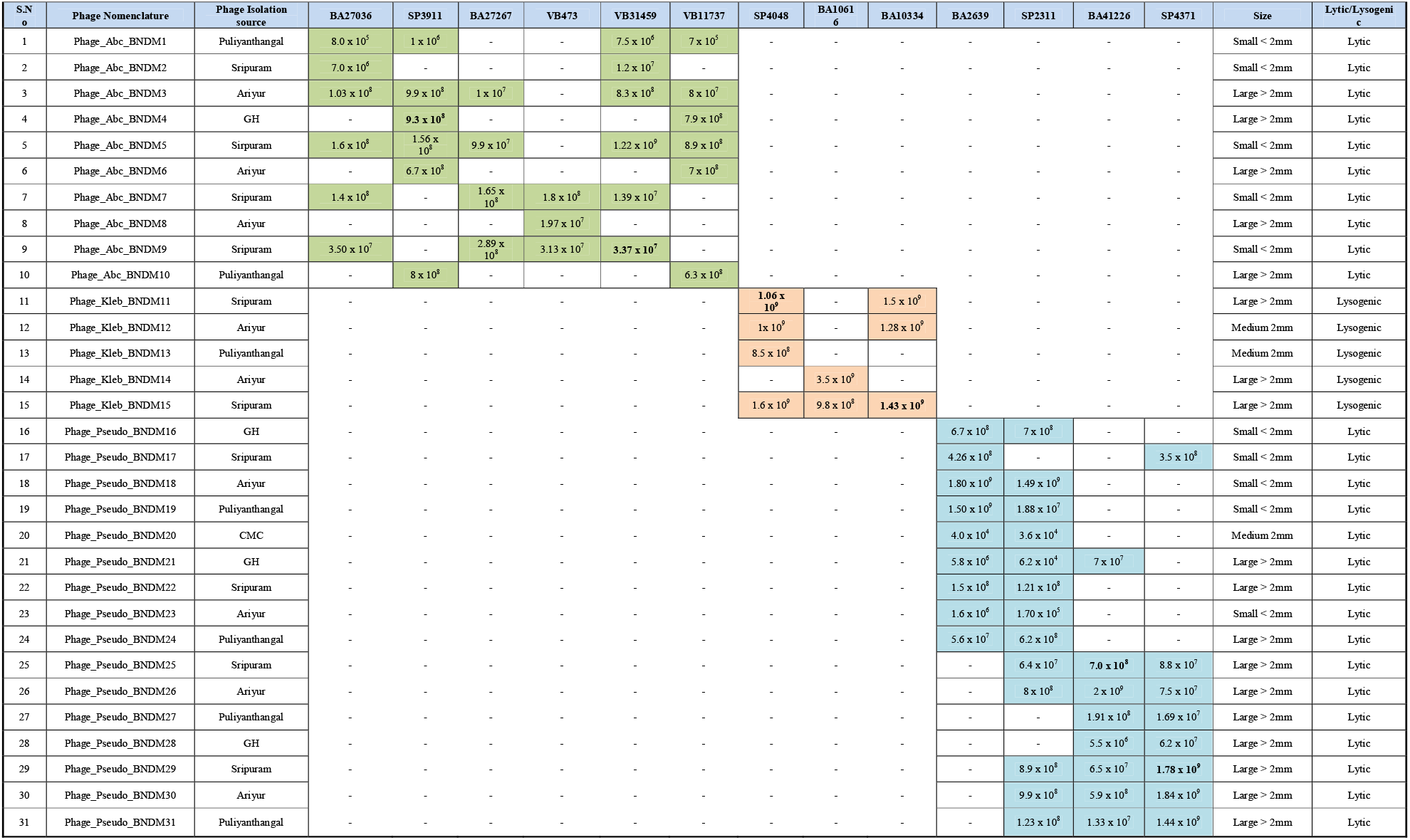
Bacteriophages tested against clinical strains and their host specificity

### Phage stability evaluation

Isolated phages were tested at a range of temperatures and pH values. The thermal stability assay showed that phages BNDM4, BNDM9, BNDM11, BNDM15, BNDM25, BNDM29 are most stable at 4 °C but activity decreased rapidly above 37 °C, with a significant reduction at 50 °C and complete inactivation at 70 °C (Figure S1A). Phages BNDM4, BNDM9, BNDM11, BNDM15, BNDM25, BNDM29 were found to be most stable at pH 6-10, indicating that neutral to slightly alkaline pH is suitable for the activity of phages (Figure S1B). Phages were all completely inactivated below pH 4 and above pH 12. For UV irradiation, phages BNDM4 and BNDM9 could resist up to 75 sec, BNDM25 and BNDM29 could resist up to 1 min in UV irradiation, and BNDM11 and BNDM15 could resist up to 2 min (Figure S1C).

### Time-dependent antimicrobial susceptibility testing of phages

Phages against *K. pneumoniae* (BNDM11, BNDM12 and BNDM13) showed inhibition of *K. pneumoniae* growth up to 6 hrs, whereas all other phages tested exhibited bacterial growth inhibition up to 12 hr. Followed by which, regrowth was observed in all wells (Table 3). Exceptionally, BNDM10 phage has shown inhibition up to 30 hrs. Interestingly, phage in combination with colistin showed complete inhibition for certain strains of *A. baumannii* and *P. aeruginosa*. Other strains of *A. baumannii* and *P. aeruginosa* showed growth inhibition up to 12 hrs. In contrast, phage with colistin combination showed antagonistic effect for two strains of *K. pneumoniae* (Table 3).

**Table 3:**
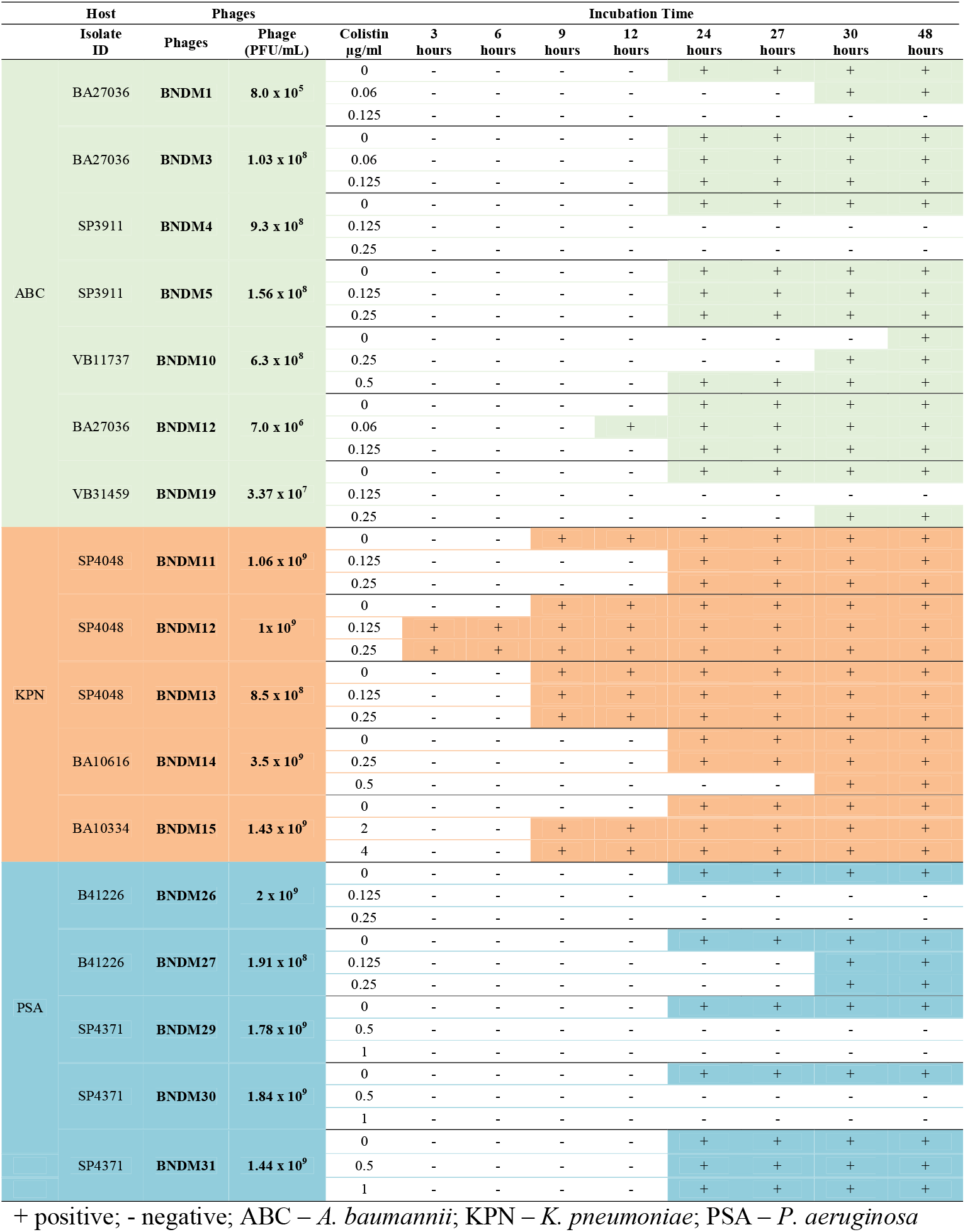
Time-dependant inhibition of bacterial cell growth by phages with and without colistin

### MBEC of colistin and phage combinations

The MBEC values for phages in combination with colistin was lower than CMT across all three species tested (Figure 1, Table 4). In *A. baumannii*, BNDM4 and BNDM9 exhibits reduced MBEC of 64 µg/ml and 0.5 µg/ml colistin concentrations in comparison to 256 µg/ml and 16 µg/ml CMT, respectively. Similarly, in *K. pneumoniae* MBECs of phage + colistin were 4 µg/ml for BNDM11 and BNDM15 phages in comparison to 32 and 16 µg/ml CMT, respectively. For *P. aeruginosa*, MBECs were 16 and 32 µg/ml for BNDM25 and BNDM29 in comparison to 64 µg/ml for CMT.

**Table 4:** MIC in comparison to MBEC of colistin with or without phages in treating ABC, KPN and PSA

| Organism | Isolate ID | Colistin MIC | Biofilm capacity | MBEC for Colistin | BNDM | Phage pfu/ml | MBEC Phage +COL (16 µg/ml) | Cocktail phage pfu/ml | Cocktail phages BNDM | MBEC for Phage cocktail + COL (16 µg/ml) |
| --- | --- | --- | --- | --- | --- | --- | --- | --- | --- | --- |
| *Acinetobacter baumannii* | SP3911 | ≤ 0.5 (I) | Strong | 256 (R) | BNDM5 | 9.3 x 10<sup>8</sup> | 64 | 1.24 x 10<sup>7</sup> | 1+3+4+5+10 | 4 (R) |
|  | VB31459 | ≤ 0.5 (I) | Strong | 16 (R) | BNDM9 | 3.37 x 10<sup>7</sup> | 512 | 5 x 10<sup>7</sup> | 1+2+3+4+9 | 0.5 (S) |
| *Pseudomonas aeruginosa* | B41226 | ≤ 0.5 (I) | Strong | 64 (R) | BNDM25 | 7.0 x 10<sup>8</sup> | 16 | 1.57 x 10<sup>7</sup> | 25+26+27+29+30+31 | 2 (I) |
|  | SP4371 | 2 (I) | Strong | 64 (R) | BNDM29 | 1.78 x 10<sup>9</sup> | 32 | 1.2 x 10<sup>7</sup> | 25+26+27+29+30+31 | 1 (I) |
| *Klebsiella pneumonia* | SP4048 | 0.5 (I) | Strong | 32 (R) | BNDM11 | 1.06 x 10<sup>9</sup> | 64 | 2.13 x 10<sup>11</sup> | 11+12+13+15 | 128 (R)/32 |
|  | BA10334 | 8 (R) | Strong | 16 (R) | BNDM15 | 1.43 x 10<sup>9</sup> | 4 | 8.4 x 10<sup>9</sup> | 11+12+15 | 512 (R) |

Isolates with Intermediate MIC but resistant to colistin in biofilm structure (MBEC); R – resistant, I – intermediate, S – susceptible; COL - colistin

**Figure 1:**
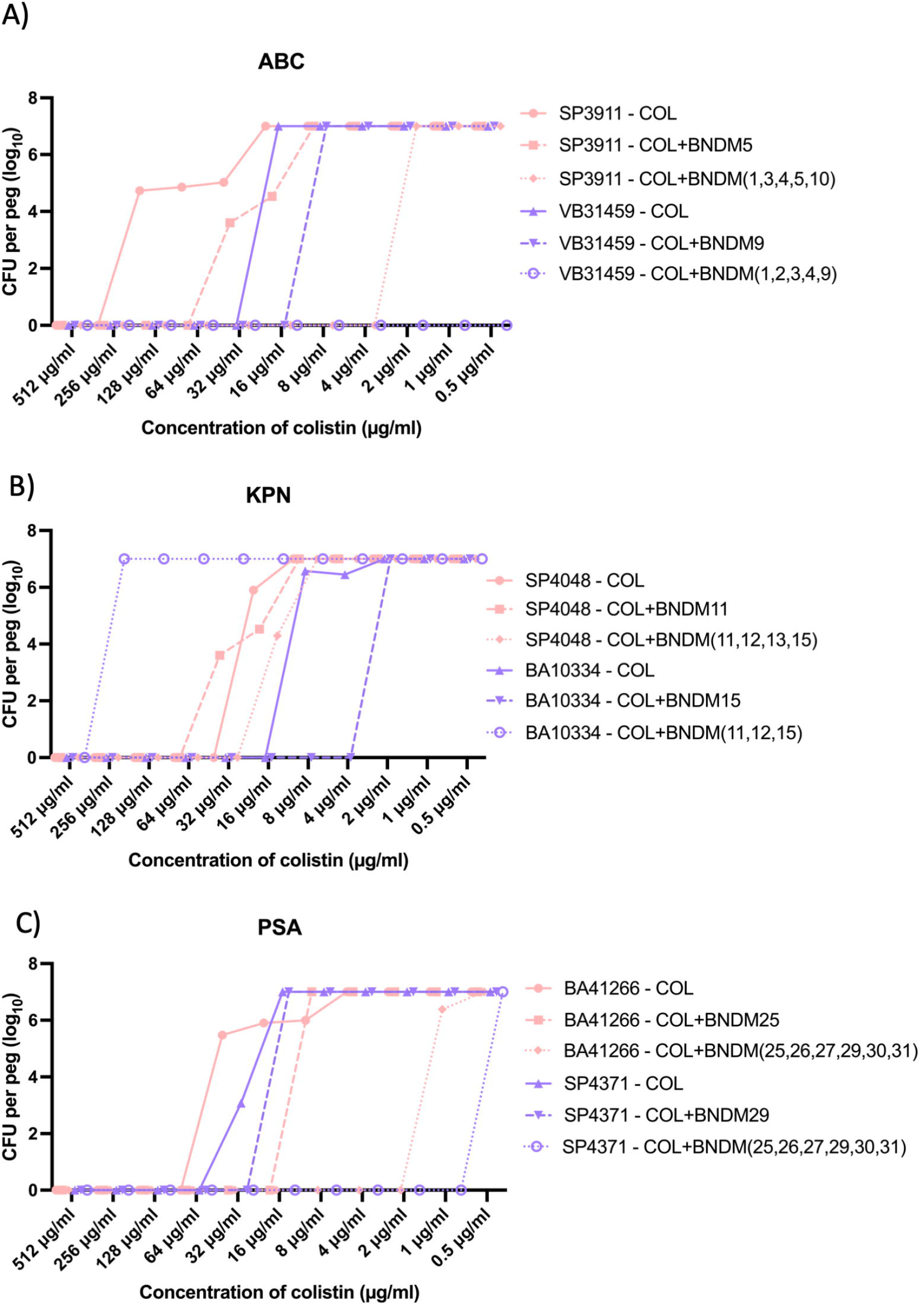
MBECs of colistin vs Phage + COL for (A) *A. baumannii* (B) *K. pneumoniae* (C) *P. aeruginosa* which shows a decreasing concentration of colistin required for clearing off biofilms.

Moreover, a cocktail of phages combined with antibiotic further reduced the colistin concentration required to eradicate biofilms (Figure 1). Interestingly, phage cocktail BNDM(1,2,3,4,9) resulted in complete eradication of biofilms even at lowest concentration of 0.5 µg/ml colistin (Figure 2). This is a 6-dilution reduction of MBEC in comparison to CMT.

**Figure 2:**
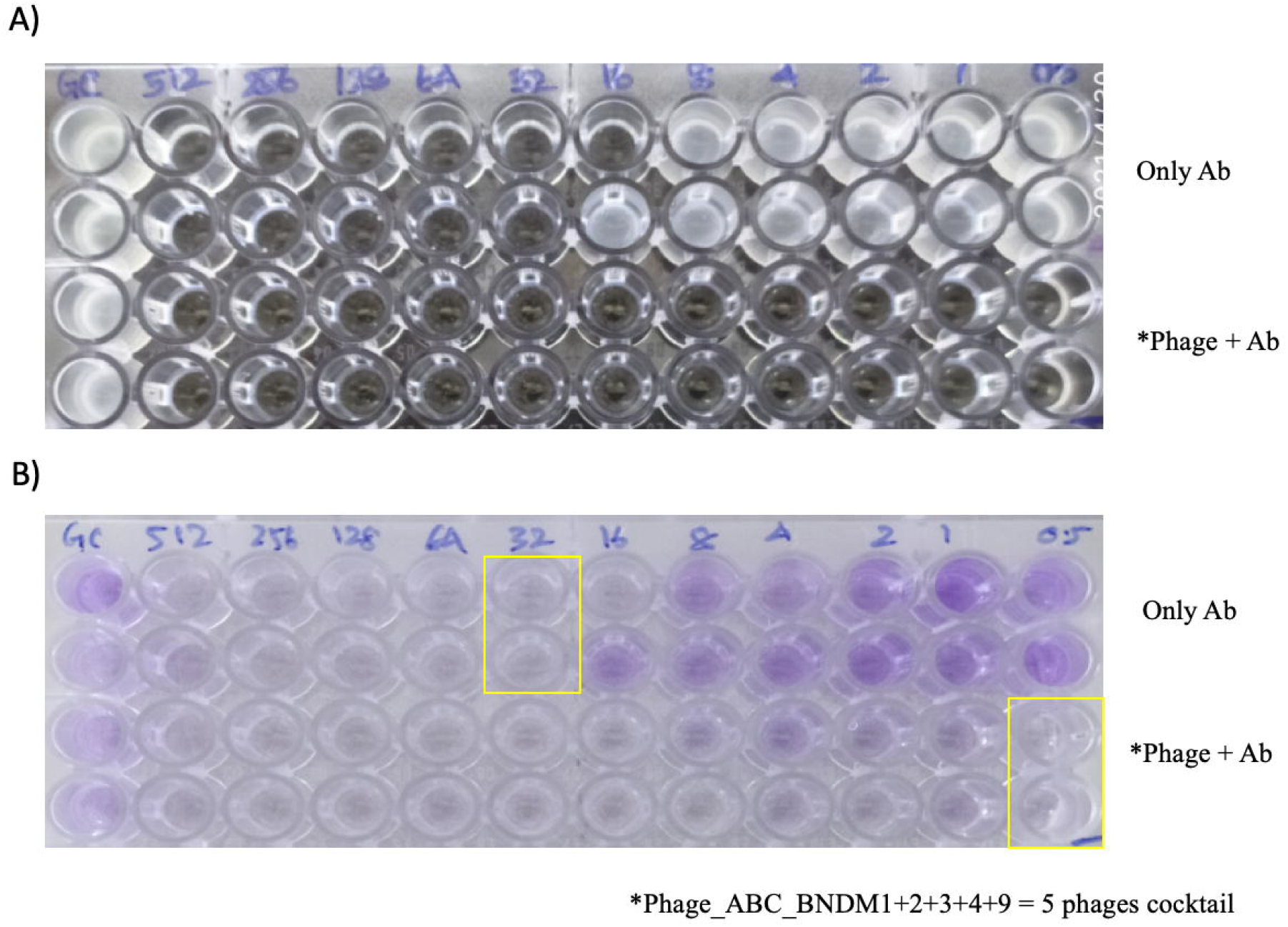
Complete eradication of VB31459 *A. baumannii* biofilms at 0.5 µg/ml colistin with phage cocktail BNDM(1,2,3,4,9) in comparison to 32 µg/ml MBEC for colistin alone as observed in (A) cells released from biofilms, and (B) biofilms on pegs stained using crystal violet.

### Qualitative estimation of biofilm eradication by colistin + phage treatment

*A. baumannii* (BNDM4) and *P. aeruginosa* (BNDM29) phages were further subjected to imaging by standard microscopy and SEM. Results revealed that phages along with sub-inhibitory concentrations of colistin (16 µg/ml) had exhibited a significant reduction in bacterial numbers and a clear eradication of biofilm extracellular matrix (ECM) in both SP3911 and SP4371 strains (Figure 3). The SEM analysis further confirms the disruption of ECM in both the strains by BNDM4 and BNDM29, respectively, in addition to shrinkage of ECM in SP4371 (Figure 4).

**Figure 3:**
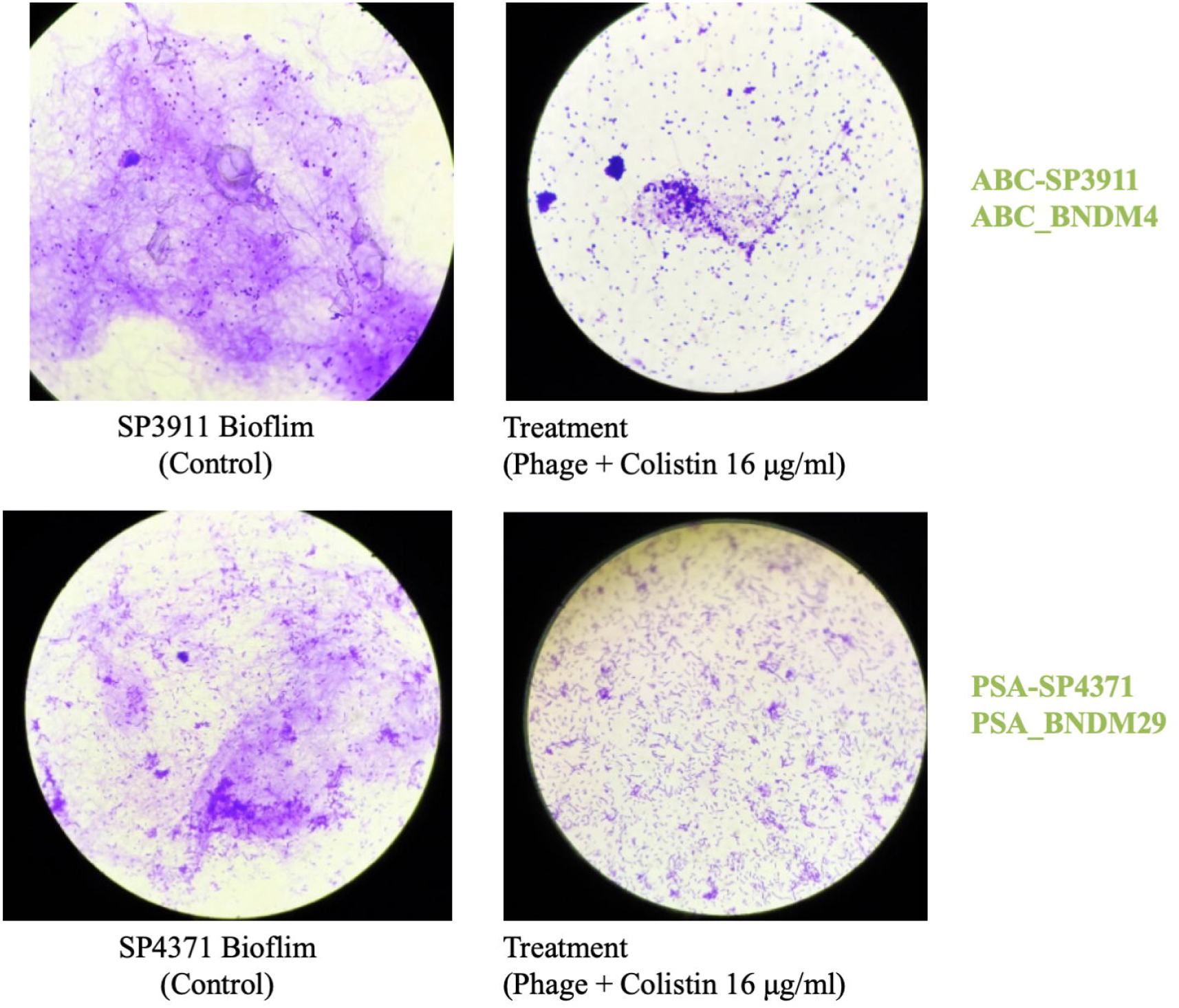
Antibiofilm efficacy of phages BNDM4 and BNDM29 depicting disruption of biofilm extracellular matrix and clumping of cells.

**Figure 4:**
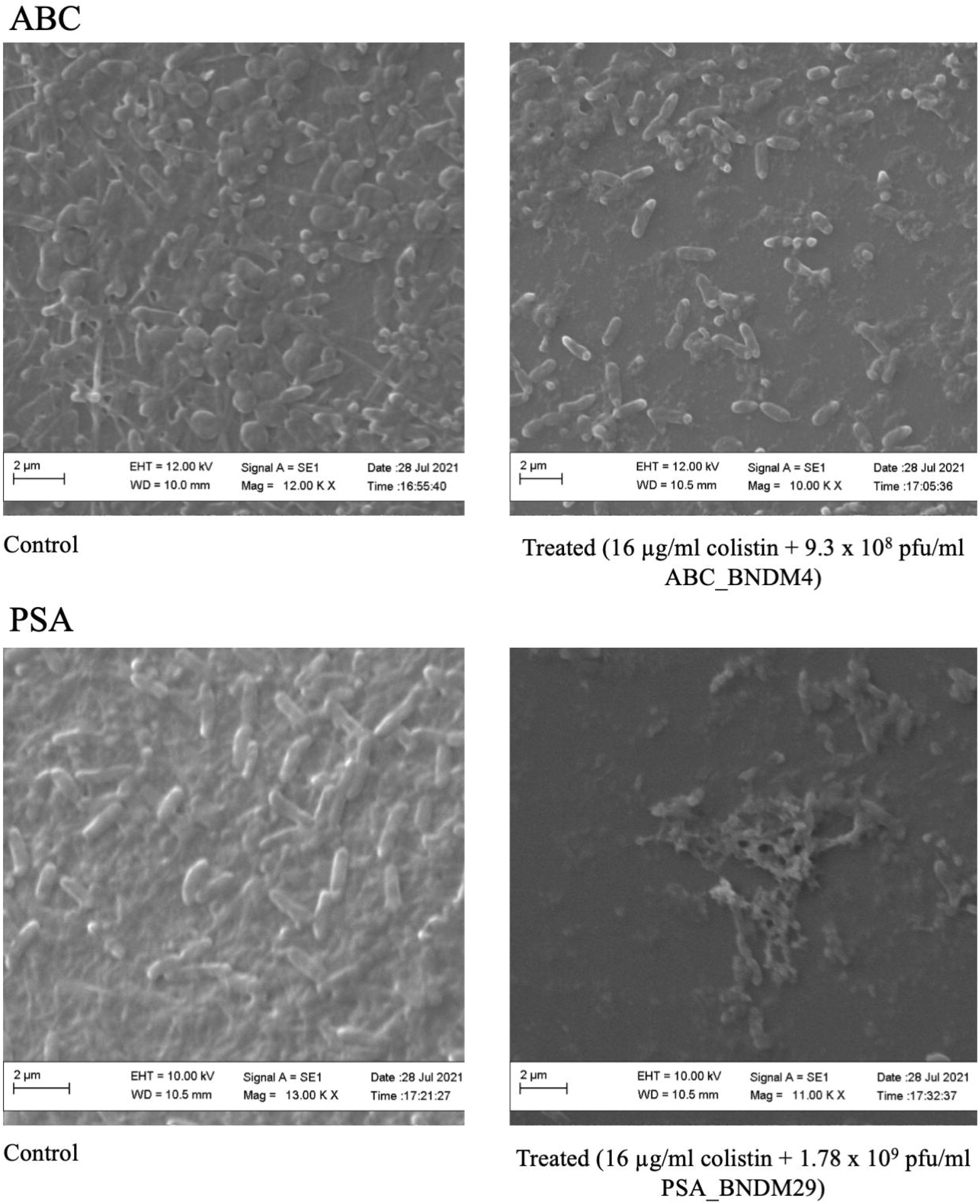
SEM imaging showing antibiofilm efficacy of Phage + COL at sub-optimal inhibitory concentrations (16 µg/ml + phages) BNDM4 and BNDM29 against A) SP3911 *A. baumannii* B) SP4371 *P. aeruginosa*, respectively.

### Host and phage genome characteristics

The clinical strains WGS revealed their genetic characteristics on AMR and virulence factors along with their prophage regions (Table 5). Novel clinical phages against specific host species were identified in this study. Genome diversity was vast in these phages even within the same host species. The phage genomes were analysed to identify their properties and safety profiles. On analysis, only *A. baumannii* phages found to have holin–endolysin complex and lytic murein transglycosylase enzymes that cleaves the peptidoglycan structures of the bacterial cell wall (BNDM2 and BNDM9) while either endolysin or holin was present in *P. aeruginosa* and *K. pneumoniae* phage sequences. Notably, no sequences associated to undesirable genes including antibiotic resistance, virulence or toxin genes were found. Few phages were found to have toxin-antitoxin system. *P. aeruginosa* phage, BNDM29 found to harbour toxic anion resistance protein. Similarly, *K. pneumoniae* phages (BNDM11 and BNDM15) found to have certain toxic protein such as HokE, SymE, RelE and TomB that destruct the host bacterial envelops.

**Table 5:**
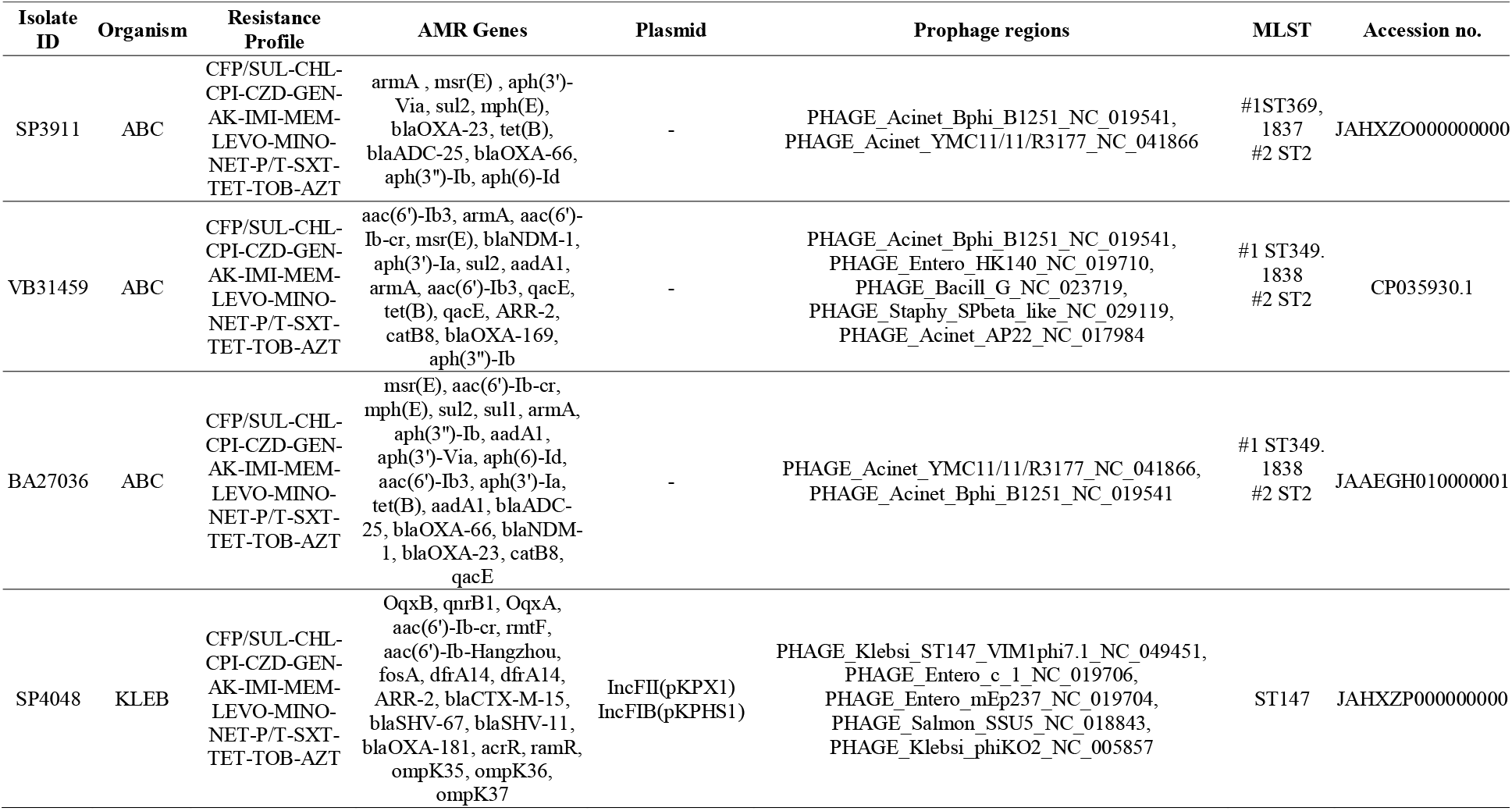
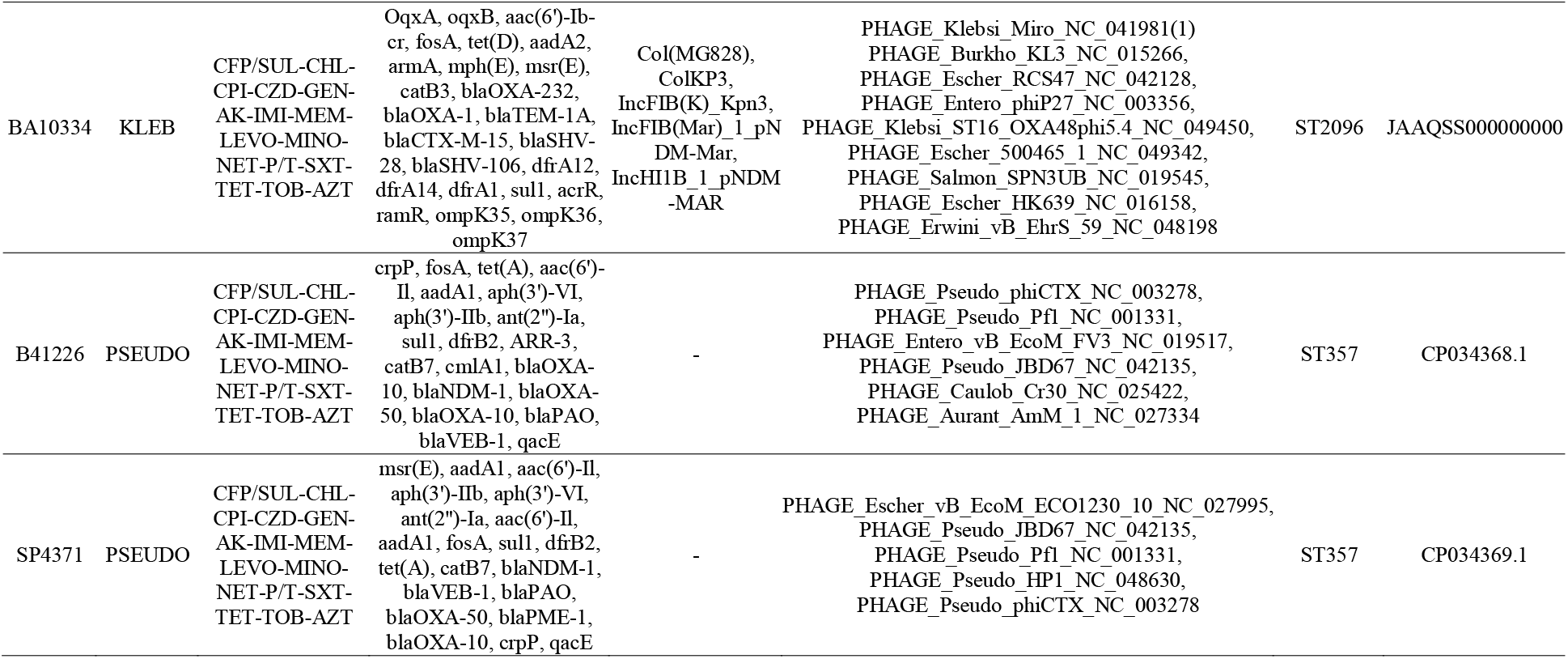
Genome characteristics of clinical host strains selected for bacteriophage treatments

In addition, the genome sequences of BNDM4, 9 and 3 against *A. baumannii*, and BNDM25 and BNDM29 for *P. aeruginosa* were compared with the global datasets from NCBI. Analysis revealed that Indian *P. aeruginosa* phages were closely related to phages from European and Chinese origin. Among *A. baumannii*, the phages were of two distinct types with no phylogenetic relation. In type I, Indian phage BNDM2 and BNDM9 were closely related to China, Israel and Thailand, while BNDM4 in type II was related to Korea and Belarus (Figure 5).

**Figure 5:**
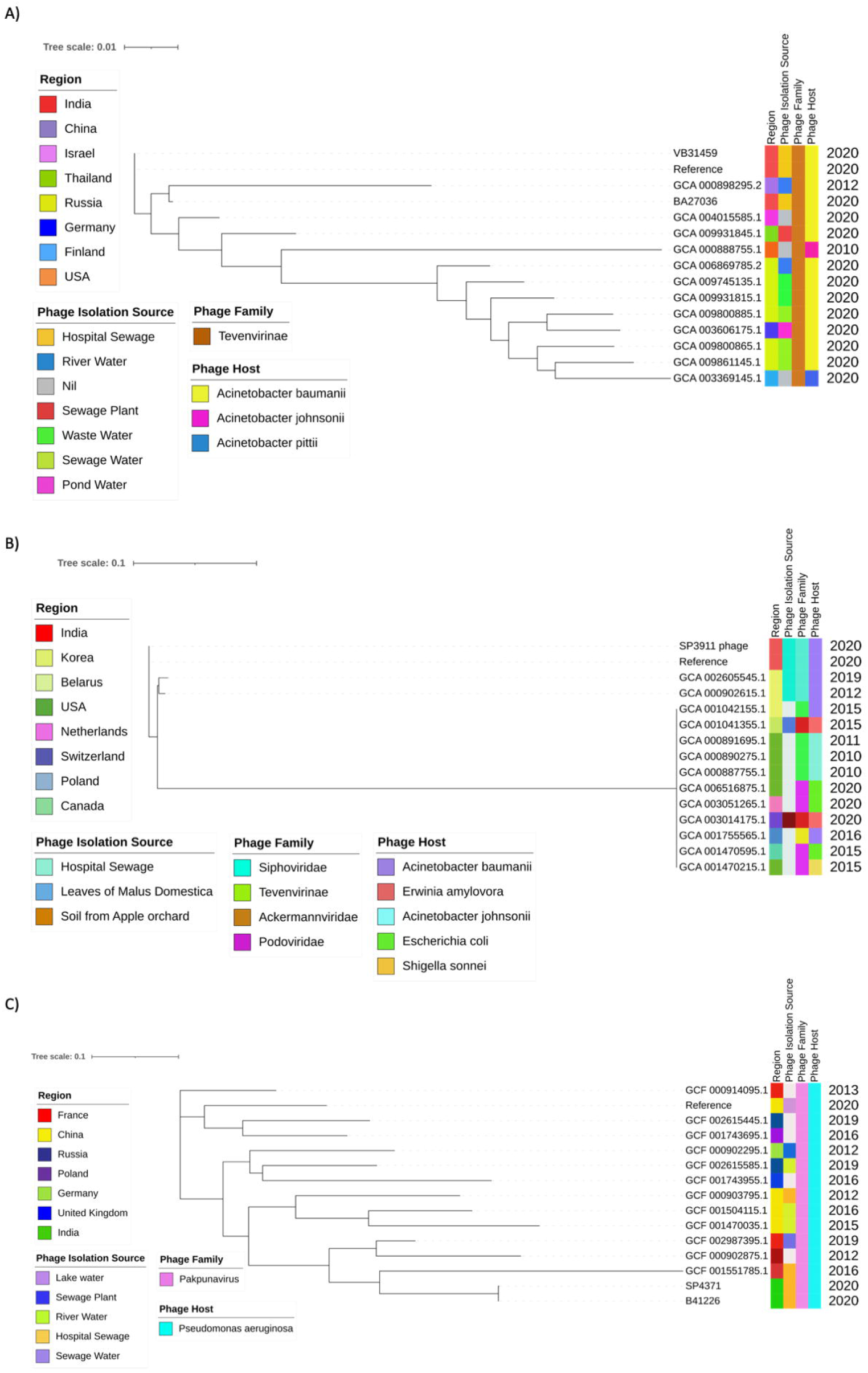
SNP based phylogeny of phages (A) for P. aeruginosa BNDM25 and BNDM29 in close relation to phages from European and Chinese origin; (B) for A. baumannii BNDM2, BNDM9, and BNDM4 with global A. baumannii phages type I Tevenvirinae and II Siphoviridae, respectively. Tevenvirinae was closely related to Finland, China, Thailand, Russia and Germany, while Siphoviridae was closely related to phages reported from Belarus.

## Discussion

Today, colistin has been the last resort or is the only effective antibiotic for the treatment of most MDR infections. Increasing resistance is being reported for such MDR infections. Most of these are nosocomial with *A. baumannii, K. pneumoniae* and *P. aeruginosa* being the commonest pathogens. Mortality due to MDR *P. aeruginosa* isolates estimated to be 18 to 61%, while for MDR and XDR *A. baumannii* is 21.2%. On the other hand, colistin resistant *K. pneumoniae*, and *A. baumannii* have been associated with high mortality range of 25 to 71%, and 28 to 70%, respectively (Mousavi et al., 2021). Moreover, biofilm formation, especially in nosocomial patients associated with medical devices, is difficult to manage to prevent recurrent infections.

In this study, the bacterial host strains selected for phage screening exhibited strong to moderate biofilm forming abilities. A high number of patients (79%) were observed to be associated with ventilation among those who expired, in comparison to 45% among discharged (P< 0.05). Ventilation is considered to be highly associated with mortality in critically ill patients (Wu et al., 2019). Patients with *A. baumannii* and *K. pneumoniae* infections were higher in the expired group, whereas *P. aeruginosa* infections were higher in discharged patients. Irrespective of outcomes, most of the isolates in these patients were strong biofilm producers.

Various strategies have been considered by researchers to overcome this challenge in critically ill patients, of which, phage therapy is widely being reconsidered as a successful alternative approach (Natasha Lipman, https://www.bbc.com/news/stories-50221375). The rationale behind this approach is mainly based on phage-antibiotic synergy; a phenomenon that is reinforced by our observations.

In this study, 31 phages specific to drug resistant *A. baumannii, K. pneumoniae* and *P. aeruginosa* were isolated from hospital sewage. Of these, 7 active phages in monotherapy exhibited time dependent inhibition of host bacteria for up to 9-12 hr following which regrowth was observed. Interestingly, colistin in combination with phages exhibited 12-24 hr growth inhibition. The MBEC of the colistin required in combination with phage has been reduced up to 16-fold. Although the effects of phages on MDR and XDR bacteria have been demonstrated earlier, only few studies have investigated the effect of phages on biofilms. This work represents one of the few studies on the effective phage against MDR/XDR biofilms synergistic with traditional antibiotics such as colistin.

Interestingly, in our study, we observed complete inhibition of a few biofilm producing *A. baumannii* strains when a phage cocktail BNDM(1,2,3,4,9) was combined with 16 µg/ml colistin. Similarly, a previous study by Wang et al. (2021) demonstrated a decrease in apparent MIC when colistin was used in combination with phage Phab24 against colistin resistant *A. baumannii*. They have also identified that phage-resistant strains displayed mutations in genes that alter the architecture of the bacterial capsule and the outer membrane using whole genome sequencing (Wang et al., 2021). Another study showed synergistic activity of some antibiotics commonly used in the treatment of urinary tract infection (UTI) with MDR *A. baumannii* phage cocktails in human urinary infection models (Bartłomiej Grygorcewicz et al,. 2021). A similar study by Jansen et al. (2018) showed that phage vB_AbaM-KARL-1, which has lytic activity against MDR *A. baumannii*, has increased antibacterial activity when combined with meropenem and colistin than with antibiotics or phages alone.

Interestingly, for *P. aeruginosa* in the planktonic stage, the phage plus colistin combination worked very well with complete inhibition in a few strains in comparison to CMT. In case of *P. aeruginosa* biofilms, BNDM25 and BNDM29 reduced the requirement of colistin 2-4 folds in combination with phages. Further, use of phage cocktail decreased the requirement of colistin up to 32-64 folds (1-2 µg/ml) in comparison to colistin monotherapy (64 µg/ml). This confirms a phage-antibiotic synergy for *P. aeruginosa* in this study. These findings are in line with the literature, where a recent study by Aghaee et al. (2021) showed that phages combined with gentamicin and ciprofloxacin combination significantly decreased the concentration of the *P. aeruginosa* by more than one log.

For *K. pneumoniae* in the planktonic stage, phages exhibited limited growth inhibition for up to 6-9 hr, while colistin combinations exhibited synergism for 2 strains and antagonism for the rest of the strains. In the biofilm stage, *K. pneumoniae* was targeted better by phage plus colistin combinations for the 2 strains, with reduction of MBECs from 16-32 to 4 µg/ml. This is supported by previous evidence where a study evaluated the therapeutic effect of a newly isolated phage (φNK5) against *K. pneumoniae* infection in an intragastric mice model and showed the characteristics of potent lytic activity (within 60 min), with a quick absorption rate (5 min) and a short latent period (10 min). Similarly, Qurat-ul-Ain et al. (2021) documented phage-cefepime and tetracycline combinations with promising therapeutic effect against MDR *K. pneumoniae*.

Phages as well as phage-derived enzymes such as endolysins and holin has also been used as a therapeutic agent for a bacterial infection as they have a lower specificity toward bacteria and able to inactivate a broader range of bacterial pathogens (Manohar et al., 2020). It is evident from our study that these enzymes were present in all three species studied. Generally, double-strand DNA (dsDNA) phages employ the holin–endolysin complex to destroy bacterial host cells (Shahin et al., 2019). Interestingly, *A. baumannii* phage genomes were predicted to encode this complex. A previous study by Briers et al. (2014) describes a group of engineered endolysin-based enzymes (Artilysins) able to penetrate the outer-membrane of Gram-negative species such as *P. aeruginosa, E. coli*, and *Salmonella* enterica. Further, no undesirable genes were identified in the phages sequenced and therefore they can be considered as safe agents for therapeutic application. Moreover, evolutionary analysis of sequenced phages with global phage sequences showed a close relationship of certain phages with the neighbouring countries.

This *in vitro* finding needs to be validated *in vivo* as the successful phage–bacteria interactions also involve other aspects such as the host immune response. Further, more investigation is necessary for dose optimisation and to establish *in vivo* antibiotic combination strategies that will ensure availability of effective formulation as a salvage therapy for the infections caused by MDR/XDR pathogens. Additionally, the development of underlying phage-resistance mechanisms needs to be elucidated since the evolution of phage resistance can be more challenging *in vivo*. Besides a potential limitation of our study is that we evaluated the synergistic effect of phages only with colistin. However, our study provides experimental data that elucidates a characteristic of phages that increased colistin susceptibility among the clinically significant pathogens such as *A. baumannii, P. aeruginosa* and *K. pneumoniae*.

## Conclusions

The study presents the efficacy of novel strain-specific phages against MDR/XDR bacterial pathogens with a complete growth inhibition of up to 6 hr. Its potential *in-vitro* activity was enhanced further with the addition of traditional antibiotic, colistin. Phage plus colistin combinations greatly reduced the MBEC of colistin up to 16 folds. Notably, a cocktail of phages exhibited supreme efficacy with complete killing at 0.5-1 µg/ml colistin concentrations. Thus, phages specific to clinical strains have a higher edge in treating nosocomial pathogens with their proven anti-biofilm efficacy. Genomic characterisation of host strains revealed prophage regions within the genome. In addition, analysis of phage genomes revealed close phylogenetic relations with phages from Europe, China and other neighbouring countries. However, phages are strain-specific phenotypically, which needs additional exploration on geographical and phylogenic matching. This study serves as a reference and can be extended to other antibiotics and phage types to assess optimum synergistic combinations to combat various drug resistant pathogens in the ongoing AMR crisis. Further, dose optimisation and antibiotic combination strategies will ensure availability of effective formulation as a salvage therapy.

## Acknowledgement

ND is a Global Challenge Fellow at The University of Sheffield, supported by a Research England QR GCRF award (10065043). The authors acknowledge The University of Sheffield, Sheffield, United Kingdom and the Christian Medical College, Vellore, India for providing basic infrastructure required for the study. The study was approved by the Institutional Review Board and Ethical Committee, Christian Medical College, Vellore, India (IRB No.: 11940 dt 27-03-2019). The authors acknowledge Vellore Institute of Technology, Vellore for facilitating SEM analysis.

---

**Figure S1:**
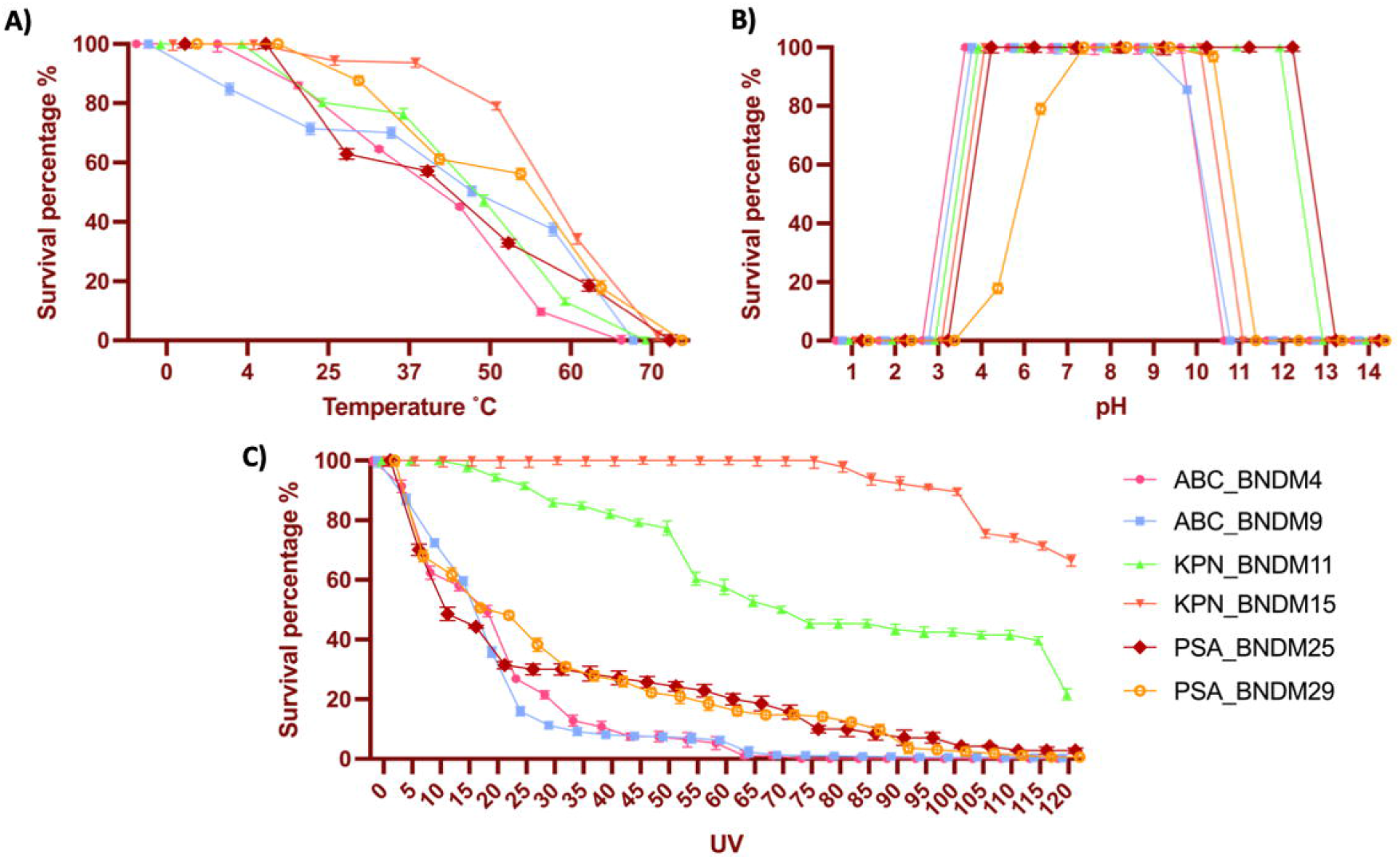
Graph exhibiting stability as survival of phages on exposure to multiple physiological factors (A) temperature, (B) pH, and (C) UV.

## Notes

### Competing Interest Statement

The authors have declared no competing interest.

